# Developmental Bioenergetic Reprogramming and Glycolytic Shift in Schizophrenia Vulnerability

**DOI:** 10.64898/2026.05.09.723970

**Authors:** J. Gilbert-Jaramillo, O. O. Folorunso, M. Castro-Guarda, T. V. Komarasamy, H. Wolosker, C. M. Palmer, Z. Sarnyai

**Affiliations:** Metabolic and Mental Health Program, McLean Hospital, Belmont, Massachusetts, USA; Stanley Center for Psychiatric Research, Broad Institute of MIT and Harvard, Cambridge, Massachusetts, USA; Center for Genomic Medicine, Massachusetts General Hospital, Boston, Massachusetts, USA; Division of Depression and Anxiety Disorders, McLean Hospital, Belmont, Massachusetts, USA; Department of Psychiatry, Harvard Medical School, Belmont, Massachusetts, USA; Basic Neuroscience Division, McLean Hospital, Belmont, Massachusetts, USA; Department of Physiology, Anatomy and Genetics, University of Oxford, United Kingdom; The Pirbright Institute, Pirbright, Woking, United Kingdom; Ruth and Bruce Rappaport Faculty of Medicine, Technion-Israel Institute of Technology, Haifa 31096, Israel; Margaret Roderick Centre for Mental Health Research, James Cook University, Townsville, QLD, Australia; Laboratory of Psychiatric Neuroscience, Australian Institute of Tropical Health and Medicine, James Cook University, Townsville, QLD, Australia; Discipline of Biomedicine and Molecular Biology, College of Medicine and Dentistry, James Cook University, Townsville, QLD, Australia; Department of Cognitive Science and Neuropsychology; University of Szeged, Szeged, Hungary

## Abstract

Schizophrenia (SZ) arises from complex gene-environment interactions, yet how early insults shape later circuit vulnerability remains unclear. Here, we investigated whether bioenergetic states represent a convergent disease signature across genetic and environmental risk factors. We analyzed transcriptional profiles across neocortical development in murine models of maternal immune activation (polyIC MIA), and serine racemase deletion (*Srr^⁻/⁻^*), extending these analyses to juvenile stages in *Srr^-/-^* and interneuron-specific NMDA receptor deletion (*Nkx2.1:Grin1^fl/fl^*), highlighting cell-type-specific metabolic vulnerability across developmental stages. In MIA, early gestation (E12.5) revealed a transient bioenergetic shift likely driven by microglial and radial glial populations, suggesting metabolic priming rather than canonical inflammatory signaling. By late gestation (E17.5), MIA induced coordinated dysregulation of neuronal glycolytic isoforms alongside mitochondrial and lipid-associated metabolic pathways, suggesting coordinated metabolic remodeling involving lipid-linked processes. In contrast, *Srr^⁻/⁻^* mice showed minimal glycolytic alterations at E17.5, indicating that isolated genetic perturbation is insufficient to recapitulate this fetal metabolic state. However, at juvenile stages, region-specific bioenergetic adaptations emerged. *Srr^⁻/⁻^* mice exhibited global cortical increases in glycolytic gene expression, with hippocampal changes potentially enriched in neuronal populations. Conversely, *Nkx2.1:Grin1^fl/fl^*interneurons showed increased glycolytic and TCA cycle transcription in the hippocampus but opposing patterns in the medial prefrontal cortex. Together, these findings identify increased glycolytic activity, potentially linked to lactate metabolism, as a partially convergent developmental mechanism bridging prenatal perturbations and later circuit dysfunction in SZ, and suggest that downstream glycolysis-linked pathways may contribute to phenotypic heterogeneity.

**Graphical Abstract:** 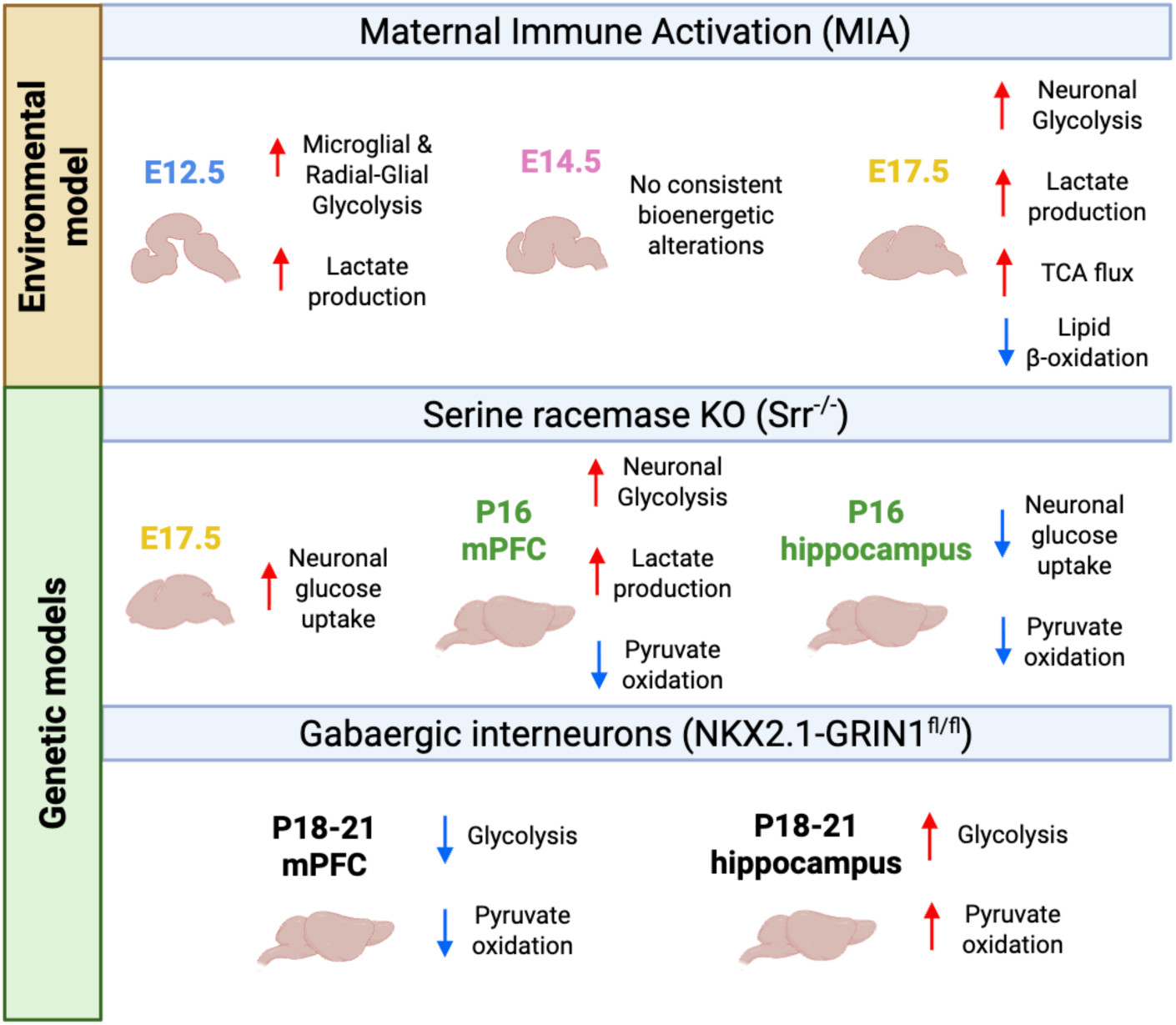

## Introduction

Psychiatric disorders such as schizophrenia (SZ) arise from complex interactions between genetic liability and environmental exposures. Genome-wide association studies have identified hundreds of SZ-associated loci implicating synaptic transmission, immune signaling, and neurodevelopmental pathways, yet no single mechanism accounts for disease onset or phenotypic heterogeneity^1^. Diverse risk factors, including polygenic variation, rare mutations, and prenatal environmental insults, may accumulate during development to shape vulnerability. Dysregulated brain bioenergetics has emerged as a plausible integrative pathway linking these multifactorial risks^1–3^.

Bioenergetic abnormalities are consistently reported in SZ. Postmortem and transcriptomic studies reveal altered expression of genes involved in glycolysis, oxidative phosphorylation, and mitochondrial function^4^, while magnetic resonance spectroscopy demonstrates increased cortical lactate and decreased ATP^5,6^. Peripheral metabolic disturbances such as insulin resistance are also observed even in antipsychotic-naïve patients^7^, suggesting that bioenergetic homeostasis may contribute to disease beyond secondary illness effects. Nevertheless, while causality remains unresolved, the early onset of bioenergetic alterations prior to major neurodevelopmental and circuit-level processes raises the possibility that they contribute to, rather than merely reflect, later dysfunction.

Current studies often rely on broad pathway enrichment analyses, which aggregate genes into generalized categories without considering tissue- or isoform-specific expression^8^. While such approaches may suggest a “Warburg-like” shift (e.g., aerobic glycolysis, an increased glycolytic ATP production in the presence of adequate oxygen supply), they rarely distinguish selective substrate reallocation, generalized upregulation, or developmental-stage–specific adaptations^9^. Single-cell and metabolic flux studies indicate glucose metabolism is ubiquitous across neurons and glia, whereas lipid metabolism is enriched in astrocytes and oligodendrocytes^10^, reflecting their roles in myelination and metabolic support. Thus, gene- and isoform-resolved analyses across defined bioenergetic modules^3^, covering glucose transport, glycolysis, lactate production, pyruvate handling, TCA cycle flux, oxidative phosphorylation, and fatty acid β-oxidation, may clarify whether dysregulation reflects coordinated reprogramming or nonspecific activation while preserving cell-type resolution.

Intrauterine brain development is a critical window during which metabolic programming may exert lasting effects^11^. Early neurogenesis relies primarily on aerobic glycolysis to sustain rapid proliferation and biosynthesis, whereas neuronal differentiation engages mitochondria with greater TCA and oxidative phosphorylation activity^12^. Glial diversification follows distinct trajectories and contributes to neuron–glia metabolic coupling through time-dependent utilization of cytosolic versus mitochondrial machinery^13,14^. Thus, disruption of these interactions may influence processes implicated in SZ, including cortical layering, neuronal migration, and synapse and circuit refinement.

Maternal immune activation (MIA) is a robust environmental risk factor for SZ. Maternal infection during pregnancy increases offspring psychosis risk^15^, which is recapitulated in preclinical rodent models using polyinosinic:polycytidylic acid (polyIC). MIA primarily acts through cytokine-mediated signaling within the maternal-fetal interface. These inflammatory processes are accompanied by alterations in oxidative stress and metabolic pathways^16^, suggesting a potential influence of prenatal immune challenges on the developmental trajectory of cellular bioenergetic states.

Genetic risk factors such as N-methyl-D-aspartate receptor (NMDAR) hypofunction are also central to SZ pathology^17^. D-serine, synthesized by serine racemase (*SRR*), acts as a key co-agonist at the NMDAR^18^. Reduced D-serine levels and altered SRR expression have been reported in SZ, and genetic variation in SRR has been associated with disease risk. In mice, *Srr* deletion lowers D-serine availability, leading to NMDAR hypofunction^19^, impaired synaptic plasticity, and cognitive deficits^20^. In contrast to high-penetrance monogenic models (e.g., *GRIN1*), SRR disruption represents a more moderate genetic liability that may better parallel the subtle effects of environmental risk factors^21^.

Herein, we hypothesize that schizophrenia risk factors converge on temporally dynamic, isoform-specific bioenergetic reorganization, and that resolving these trajectories may distinguish primary metabolic vulnerabilities from compensatory adaptations, thereby advancing bioenergetics as a framework for causal inference and patient stratification. To assess this, we structured our analyses along a developmental hierarchy spanning embryonic, juvenile, and cell-type–resolved dimensions. Using a curated, tissue- and isoform-resolved framework, modified from Sahay et al., 2024^3^, we examined bioenergetic genes across key modules, including glucose transport, aerobic glycolysis, pyruvate metabolism, TCA flux, and lipid β-oxidation, to uncover selective signatures potentially masked in generalized analyses. At the embryonic stage, we first interrogated environmental risk conditions using maternal immune activation (MIA)^22^ and complemented in our *Srr^−/−^*model to capture convergent genetic perturbation of early bioenergetic programs within the neocortex. We then extended analyses of the *Srr^−/−^* model to the medial prefrontal cortex (mPFC) during juvenile periods, a stage characterized by emergent synaptic maturation where NMDA signaling is pivotal. We next compared this to the hippocampus another area of importance in SZ yet with a different developmental trait. Lastly, using publicly available datasets^23^, we examined bioenergetic changes in a model of NMDAr deletion within inhibitory neuronal progenitors (*NKX2.1:Grin1^fl/fl^*); cells that populate both the mPFC and the hippocampus.

## Methods

### Mice

#### Serine racemase knockout (*Srr*^-/-^ mice)

C57BL6/J wild-type (Jackson Lab) and heterozygous (SR^+/-^)^24^ were used for breeding. For embryonic experiments, SR^+/-^ female mice were timed mated with either SR^+/-^ or WT males. Timed mating was initiated in the evening, and female mice were checked for vaginal plugs the following morning. Embryonic day 0 (E0) was the first day a female presented with a vaginal plug. Pregnant female mice were group-housed until E14, then single-housed until embryo extraction. For post-natal experiments, SR^+/-^females were left with SR^+/-^ or WT males for one week after initial pairing, group-housed with other pregnant dams until two weeks after initial pairing, and then single-housed until the time of pup collection. Mice were maintained in polycarbonate cages on a 12-hour light/dark cycle and given access to food and water *ad libitum*.

#### Embryonic Brains

Brains were extracted at embryonic day 17. Dams were anesthetized with an intraperitoneal injection of Ketamine (100mg/kg)/Xylazine (5mg/kg) toe pinched for non-reflex. before extraction. Embryonic brains were then extracted in 1xPBS using a dissection microscope, excess PBS removed and immediately frozen on dry ice and stored at -80°C. All embryos were genotyped, and both male and female brains were used for embryonic experiments as reported elsewhere^25^.

#### Post-natal Brains

At post-natal day (PND) 16, male mice were deeply anesthetized with isoflurane administered in a small chamber. The medial prefrontal cortex and hippocampus were dissected out and then immediately frozen on dry ice and stored at -80°C^26^.

#### Maternal Immune Activation (MIA) mice

<u>P</u>reviously published datasets were used in this study. Briefly, timed-pregnant C57BL/6N dams were subjected to maternal immune activation (MIA) via poly(I:C) challenge at E12.5. To ensure robust immune responsiveness and minimize inter-subject variability, dams were pre-screened; those exhibiting immune reactivity in the lowest 25th percentile (based on serum IL-6 levels measured 2.5 hours post-injection) were excluded from the study. Cortical tissues (dorsal telencephalon/pallium) were collected from offspring across four developmental time points: E12.5 (+6 hr), E14.5, E17.5, and P0^22,27^.

#### NKX2.1:Grin1^fl/fl^ mice

Previously published datasets were were used in this study. Briefly, datasets were generated from single-cell RNA sequencing of juvenile brain tissues (P18–P20) from pups from breed of conditional Grin1 floxed mice with Nkx2.1-Cre driver lines to achieve interneuron-specific deletion, as described^23^.

### Generation and culture of human cortical progenitors

Culture and differentiation of human pluripotent stem cells (iPSCs) was carried out as previously published under the same Ethical approval. Cell lines included SFC841-03-01, SFC-840-03-03, and SFC856-03-04^28^. Briefly, ∼95% confluent iPSC cultures were neuralized over 6-7 days neuronal induction media [NIM: NMM, 100 mM LDN193189, 10 µM SB431542] with daily feedings. On day 6-7, cells were washed once with PBS and incubated with 0.5 mM EDTA in PBS for 5 min at 37°C, 5% carbon dioxide. Cells were pelleted at 300 rcf for 3 min and resuspended, as clumps, in neuronal maintenance media [NMM: 50% Neurobasal medium, 50% DMEMF:F12 medium, 2 mM Glutamax, 1X B-27 Supplement, 1X N-2Supplement] supplemented with 10 µM ROCKi. After 5 days, cells were replated (split 1:2) as detailed before. Following three extra days in culture, cells were either replated (split 1:2) or stored in LN2 (early first trimester NPCs). Two additional passages, each of 3 days, were further conducted. Cells from the final passage were replated as single cells suspension (late first / mid second trimester NPCs). For viral infection, cells were incubated for 2 h at 37°C, 5% carbon dioxide with media-containing ZIKV MP1751 at an M.O.I. of 1^28,29^. Media-containing virus was then removed, and cells were washed once with PBS before adding fresh NMM. After 24 h, cells were washed once with PBS and collected with RLT+ (Qiagen - RNeasy Mini Kit Plus) and further stored at -80°C until processing.

### RNA extraction and Quantitative polymerase chain reaction

RNA from embryonic and postnatal mouse tissue and/or cell pellets was isolated following the manufacturer’s procedure (Qiagen RNeasy Mini Kit; Cat. No. 74104 and 74106). A DNAse step was conducted for all the samples to ensure RNA purity and RNA concentration was quantified using a Nanodrop. RNA was reversed transcribed into complementary DNA (cDNA) using High-Capacity RNA-to-cDNA™ or SuperScript^TM^ IV Reverse Transcriptase Kit. Oligonucleotide sequences were obtained from PrimerBank (Supplementary Table 1). Gene expression was determined using the double stranded DNA binding dye SYBR™ Green. qPCR reactions comprised 3 ριg cDNA per reaction, in triplicates over 40 cycles. The threshold cycle (2−ΔΔCt) method of comparative PCR was used to analyse the results. 2−ΔCt was calculated by the normalization of the sample Ct value to the average Ct value of two housekeeper genes, β2M and HPRT1.

### Public dataset reanalysis and bioenergetic gene set scoring

Publicly available transcriptomic datasets were reanalysed to investigate bioenergetic transcriptional programs. Bulk RNA-sequencing (GSE166376)^22^ and single-cell RNA-sequencing (GSE156201)^23^ data were obtained from the Gene Expression Omnibus (GEO). Bulk RNA-sequencing differential expression analysis and normalisation of gene counts were carried out using the DESeq2 pipeline^30^. Genes with fewer than ten counts in at least the minimum group size of samples in either condition were removed from the analysis as lowly expressed. For visualisation, exploratory analysis, and mean-variance stabilisation, counts were normalised using the variance stabilising transformation (VST)^31^. Differential gene expression between groups was assessed from raw counts, with Log2fold changes refined using the apeglm shrinkage method to ensure accurate group comparisons^32^. Heatmaps and box plots were generated using the ggplot2^33^, ComplexHeatmap^34,35^, ggpubr^36^, and/or dplyr^37^ packages. Single-cell RNA-seq data were analysed using Seurat package^38–41^. Low-quality cells were excluded (<200 or >6000 genes, <500 UMIs, >10% mitochondrial reads). Data were log-normalised, and the 3,000 most variable features were identified. Following scaling and regression of mitochondrial content, dimensionality reduction was performed using PCA and UMAP. Metabolic pathway activity was quantified per cell using AddModuleScore. All analyses and coding were conducted in R v4.4.3 and RStudio.

To assess pathway-level alterations, curated gene sets representing key metabolic processes (including glycolysis, lactate metabolism, pyruvate metabolism, TCA cycle activity, and mitochondrial fatty acid oxidation) were defined (Supplementary Table 1). Gene selection was based on key referenced genes in the literature for each pathway as well as mouse ortholog genes to humans with their respective cell-type-specific isoform where applicable. Following a similar approach to Ling et al. (2024)^42^, gene set expression scores were calculated by aggregating normalized expression values across genes within each pathway. In bulk RNA-seq, scores were computed per sample and compared across groups. In single-cell RNA-seq, scores were calculated per cell within each brain region. These bioenergetic expression signatures capture coordinated transcriptional activity across bioenergetic pathways, enabling comparison of metabolic states across conditions and cell populations.

### Statistical analysis and software

For qPCR results, where normal distribution was met, parametric tests were performed. In cases of missing values and/or values of zero, we performed mixed-effects model with Holm-Šidák correction. Multiple t-tests were used to analyse datasets from human cortical progenitors. A calculated P-value less than 0.05 was reported as significantly different. All the statistical tests were performed using GraphPad Prism 10 v10.6.1. For RNAseq statistical significance was defined as a Benjamini-Hochberg adjusted p-value < 0.05 conducted in R v4.4.3 and RStudio. Representation schemes were created with BioRender.com.

## Results

### Early glycolytic transcriptional responses in the neocortex following MIA are cell-type specific

Given the limited characterization of intrauterine cortical responses in SZ risk models, we examined whether bioenergetic transcriptional disturbances reported in the adult brain emerge during intrauterine life in environmental and genetic risk models. To determine whether maternal immune activation (MIA) induces acute metabolic transcriptional responses over the normal transcriptional programs in the developing cortex^43^, we analyzed gene expression in the developing neocortex at embryonic day (E)12.5, 6 h post-maternal exposure to poly:IC (MIA). Analysis revealed increased expression of multiple genes involved in glucose entry, glycolysis, and lactate production including *Slc2a1, Slc2a3, Slc2a4, Gapdh*, *Aldoa, Pgk1, Tpi1, Hk2, Pdk1*, and *Ldha* (Figure 1A). In contrast, little to no changes were observed in genes associated with mitochondrial oxidative metabolism and fatty acid oxidation (Figure 1B). Isoform-level analysis, informed by cell-type expression profiles, indicated that several upregulated transcripts, including *Hk2* and *Pdk1*, are predominantly expressed in glial cells including radial-glia and microglia.

**Figure 1.**
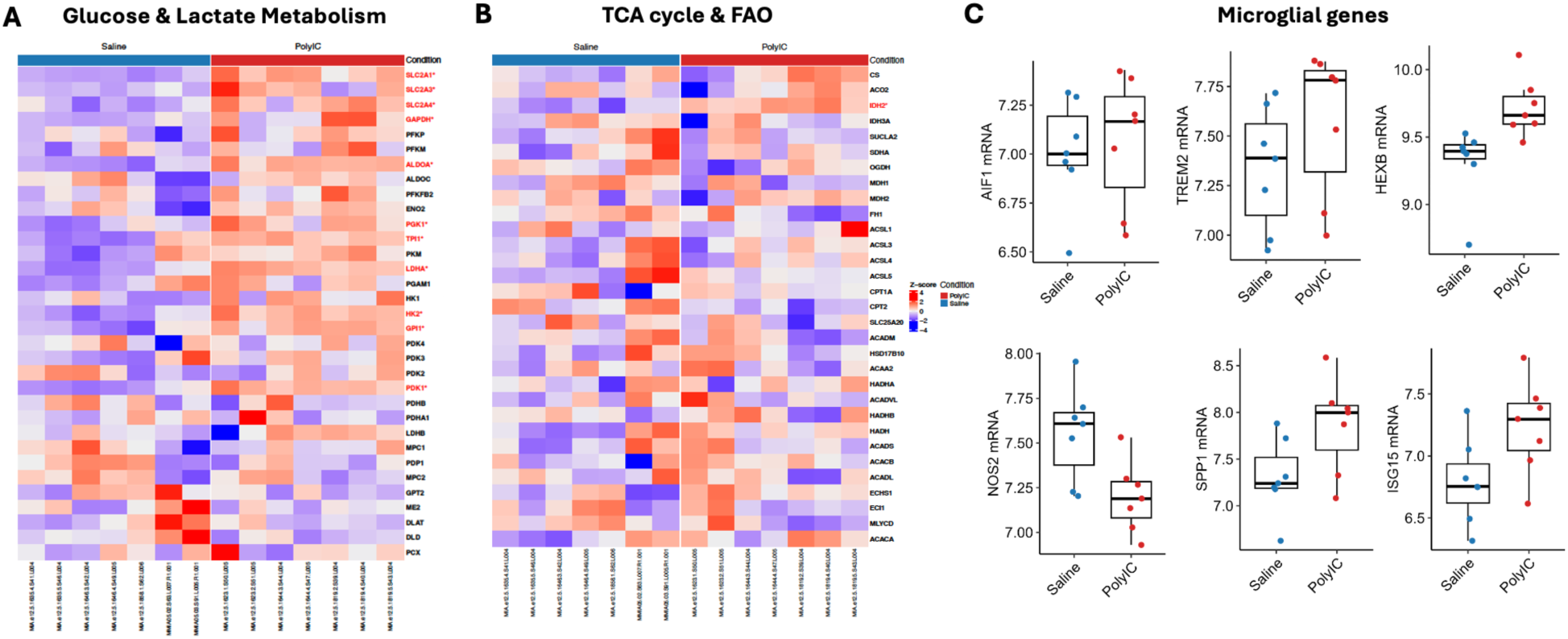
MIA induces lactate glycolysis reprogramming in the developmental embryonic cortex. Heatmaps showing the p-adjusted expression values for multiple genes involved in glycolysis, lactate production (A), pyruvate metabolism, TCA cycle flux and fatty acid β-oxidation (B). Red showing increase in mRNA expression compared to untreated, blue showing downregulation. Significance is highlighted by an *. (C) Box plots of the normalised counts display the median (interquartile range) of multiple microglial-specific genes across the MIA and saline groups. Results correspond to bulk RNA-seq neocortex of E12.5.

To determine whether these transcriptional changes reflected global alterations or cell-type–specific responses, we assessed the expression of genes associated with microglial abundance and activation. Across markers of homeostatic identity (*Tmem119, P2ry12, Sall1, Fcrls, Cx3cr1, Hexb, Csf1r, Aif1*), surveillance (*Trem2, Cd68*), pro-inflammatory signaling (*Il18, Nos2, Fcgr1, Fcgr3, Lyz2, Spp1, ApoE, Lpl, Mki67, Pycard, Ifitm3, Irf7, Isg15*), and anti-inflammatory regulation (*Tgfb1, Mrc1, Igf1*)^44^, no consistent transcriptional changes indicative of altered microglial abundance or classical activation states were detected following MIA. Selected genes with the largest median differences are shown in Figure 1C, with the remaining panel provided in Supplementary Figure 3.

Because classical inflammatory mediators were largely absent or below detection thresholds (Supplementary Table 2), we next examined whether cell-type composition could obscure immune-related signals in bulk RNA sequencing^8,10^. Specifically, the presence of neuronal progenitor populations with intrinsically low inflammatory responsiveness^45^ may mask transcriptional changes arising from immune-competent cells. To address this, given that both poly(I:C) and neurotropic Zika virus infection engage TLR3 signaling^46^, we exposed human pluripotent stem cell-derived cortical progenitor pools at different stages of differentiation to virus^28,29^ and assessed the expression of key innate immune genes. In early-stage progenitor populations resembling first-trimester forebrain development, IRF3 expression was significantly reduced (Supplementary Figure 4A). In more developmentally advanced populations, *IRF3* remained reduced (not significant), whereas *CCL4, CCL5, IL1B, TNFA*, and *MMP9* were significantly increased (Supplementary Figure 4B). Together, these findings are consistent with the possibility that the predominance of early cortical progenitors with limited innate immune responsiveness may attenuate inflammatory transcriptional signatures detected in bulk RNA sequencing datasets.

### Late gestation reveals neuronal glycolytic transcriptional upregulation in MIA models

To determine whether early transcriptional alterations persisted during subsequent stages of cortical development, we analyzed gene expression of bioenergetic genes at E14.5, a period of active neurogenesis. At this stage, few significant changes were observed across all the assessed pathways with the most consistent coordinated pathway-level regulation displaying *Pfkp* expression significantly increased, and *Ldha, Ldhb,* and *Pdk3* significantly decreased (Figure 2A). We next examined transcriptional changes at E17.5, corresponding to a later stage of cortical maturation. At this timepoint, multiple genes were significantly upregulated (Figure 2B). Analysis of genes involved in pyruvate metabolism and mitochondrial import, showed no significant changes at E14.5 whereas *Pdha1, Pcp1,* and *Mpc2*, showed significant increments at E17.5. Isoform analysis revealed cell-type–associated shifts, with reduced expression of glial-enriched isoforms (*Slc2a1, Slc2a4, Aldoc, Hk2, Pdk2*) and increased expression of neuronal isoforms (*Slc2a3, Eno2, Hk1*).

**Figure 2.**
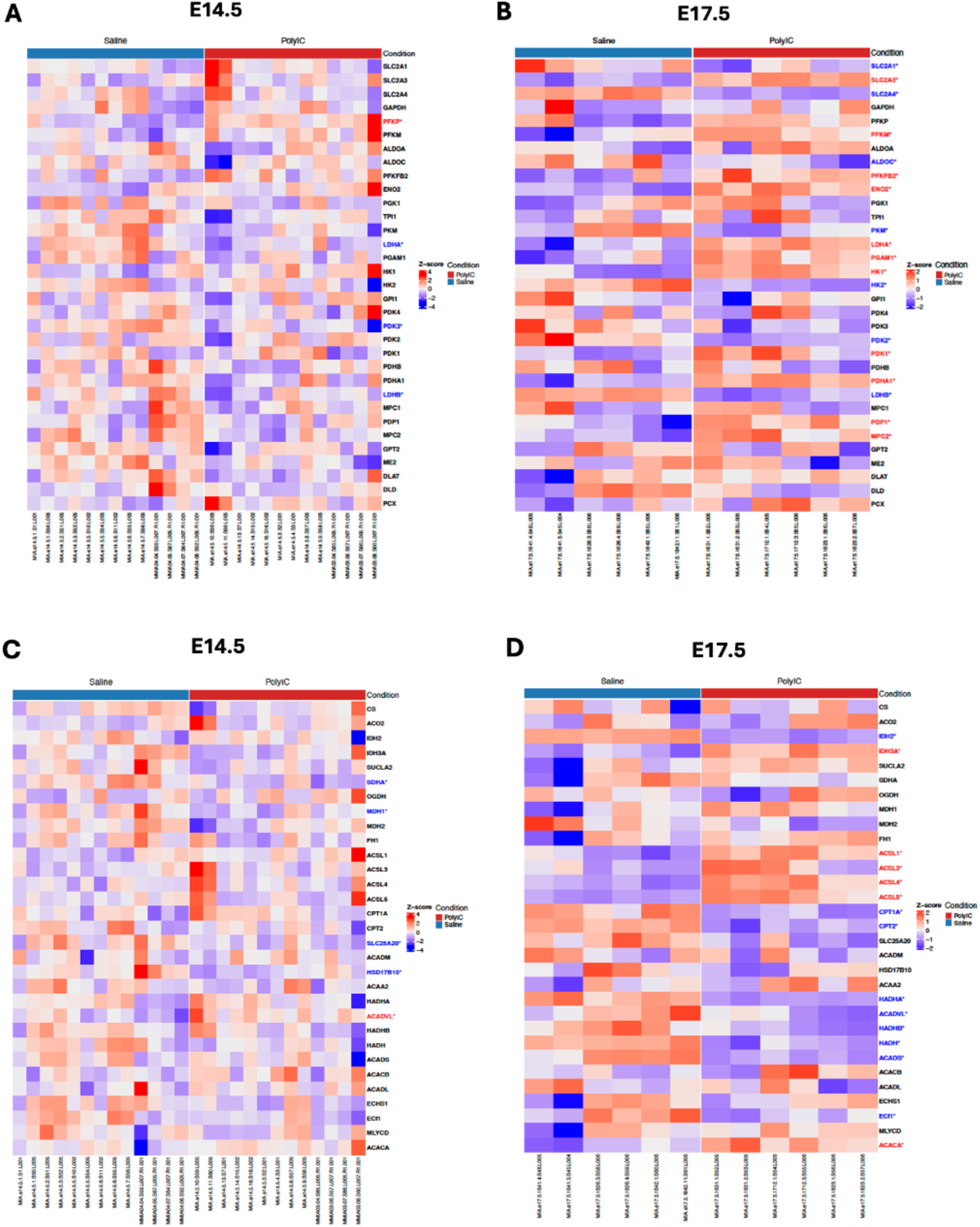
MIA induces aerobic glycolytic preference and lipid remodeling during late gestation. Heatmaps showing the p-adjusted expression values for multiple genes involved in glycolysis and lactate production at (A) E14.5, and (B) E17.5, as well as pyruvate metabolism, TCA cycle flux and fatty acid oxidation (C) E14.5, and (D) E17.5. Red showing increase in mRNA expression compared to untreated, blue showing downregulation. Significance is highlighted by an *. Results correspond to bulk RNA-seq.

To assess whether additional bioenergetic pathways were affected, we examined genes associated with the tricarboxylic acid (TCA) cycle and mitochondrial fatty acid oxidation. At E14.5 (Figure 2C), few significant changes were detected. In contrast, at E17.5 (Figure 2D), isoform-specific alterations were observed in TCA cycle–related genes, including decreased expression of the astroglial isoform *Idh2* and increased expression of the neuronal isoform *Idh3a*. Similarly, multiple genes involved in mitochondrial β-oxidation were significantly reduced, including those mediating fatty acid transport (*Cpt1a, Cpt2*), acyl-CoA dehydrogenation (*Acads, Acadvl*), auxiliary oxidation (*Eci1*), mitochondrial fatty acid metabolism (*Hadh*), components of the mitochondrial trifunctional complex (*Hadha, Hadhb*), fatty acid activation (*Acaca*), and long-chain acyl-CoA synthetases (*Acsl1, Acsl3, Acsl4, Acsl5*).

To assess whether similar transcriptional changes are present in a genetic risk model, we analyzed embryos lacking serine racemase (*Srr^−/−^*), a model of NMDA receptor hypofunction^18,19^. Gene expression was examined at E17.5, corresponding to the stage showing the largest transcriptional changes in the MIA model (Figure 3). Quantitative PCR revealed no significant changes in the expression of genes involved in glucose uptake, metabolism, and oxidation, as well as pyruvate metabolism and TCA flux.

**Figure 3.**
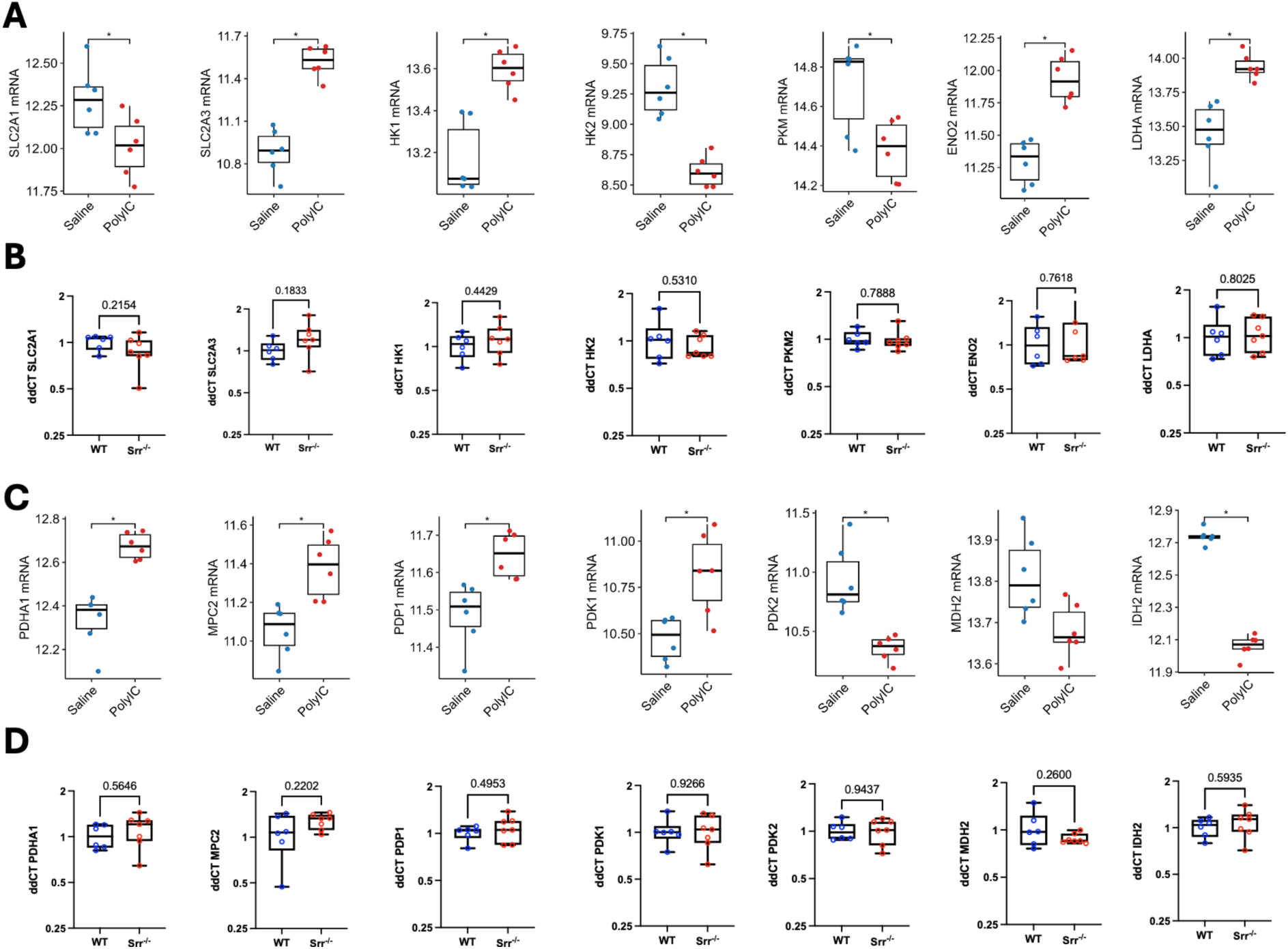
*Srr^-/-^* does not induce glycolytic transcriptional increases during late gestion. Box plots showing the mRNA expression levels of (A-B) glycolytic genes, and (C-D) genes related to pyruvate metabolism and TCA cycle flux at E17.5. Panels A and C correspond to the MIA model whilst B and D correspond to *Srr^-/-^* model. Blue and red dots represent control and experimental mice, respectively. For the *Srr^-/-^*model, statistical analysis was performed on relative expression values from seven knockout (KO) and six wildtype (WT) mice. Box plots (panels A and C) show median (interquartile range). Statistical significance (adjusted p-values) was determined using DESeq20. Box plots (B and D) show median (min to max values). Data were analyzed using a t-test with Welch’s t-test correction following log transformation to account for log-normal distribution. Significance is displayed for * p ≥ 0.05.

### Postnatal glycolytic transcriptional changes exhibit region- and cell-type specificity in a genetic SZ risk model

As the embryonic signature between the MIA and the *Srr^-/-^*may reflect differential transcriptional rewiring and a bioenergetic diversion between risk models of schizophrenia, examined expression of the same metabolic gene panel in postnatal day (P)16 *Srr^−/−^* mice. Because this period englobes critical milestones for cortical and hippocampal maturation, we focused our analysis on the medial prefrontal cortex (mPFC) and hippocampus^47^. At P16, mPFC and hippocampus exhibited significant bioenergetic reprogramming, though the nature of these adaptations varied by region. The mPFC displayed a broad transcriptional upregulation of glucose entry receptors encompassing both *Slc2a1* (primarily glial/endothelial) and *Slc2a3* (neuronal) isoforms^48^, as well as elevated genes involved in glycolysis (*Hk1*, *Hk2*, *Pkm2*, *Eno2*, *Ldha*), pyruvate metabolism (*Pdha1, Mpc2, Pdk1*) and TCA flux (*Idh2*) (Figure 4A–B). In contrast, despite the resemblance in the elevation of glycolytic genes (*Hk1*, *Hk2*, *Pkm2*, and *Ldha*) and TCA flux via *Idh2*, the hippocampus showed a neuronal restricted profile for glucose entry (*Slc2a3*) and a distinct pyruvate metabolism profile (*Pdha1*, *Pdp1*) (Figure 4C–D).

**Figure 4.**
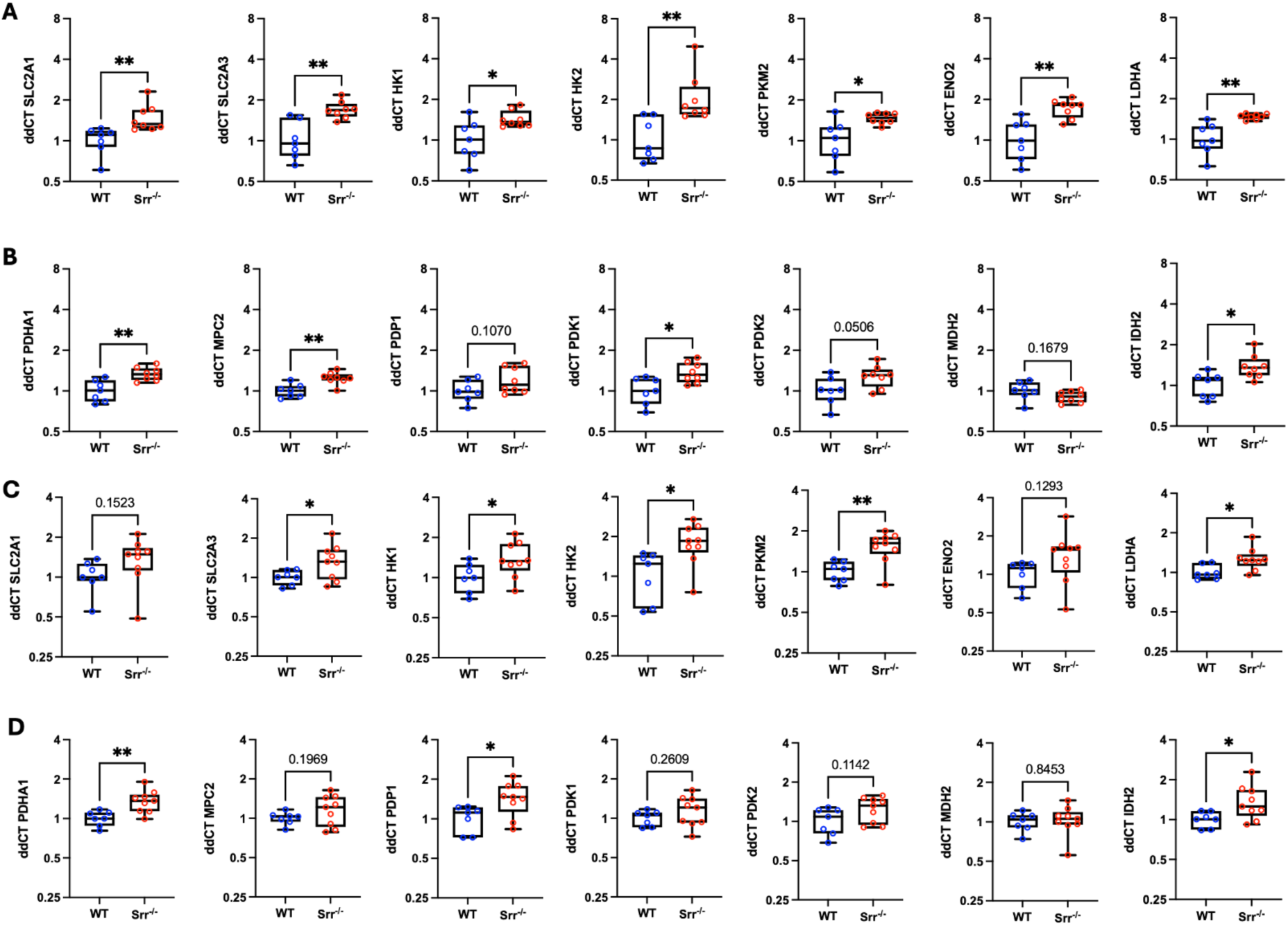
Juvenile *Srr^-/-^* mice show distinct bioenergetic profiles across different brain regions. Box plots displaying mRNA expression levels in the *Srr^⁻/⁻^* model at P16. (A-C) Glycolytic genes and (B-D) genes related to pyruvate metabolism and TCA cycle flux. Panels A and B correspond to medial prefrontal cortex (mPFC), while panels C and D correspond to hippocampus. Blue and red dots represent wildtype (WT) and knockout (KO) mice, respectively. Bos show median (min to max values). Statistical analysis was performed on relative mRNA expression values from seven KO and six WT mice using a t-test with Welch’s t-test correction following log transformation to account for log-normal distribution. Plots display mean ± SD. Significance is displayed for * p ≥ 0.05, ** p ≥ 0.005, and *** p ≥ 0.0005.

This regional divergence raised the possibility that specific cell populations contribute differentially to these bioenergetic phenotypes. Given the high metabolic demand of inhibitory circuits^49^ and their established role in schizophrenia pathology^50^, we sought to determine whether interneuron-specific bioenergetic transcriptional dysregulation could explain the region-specific patterns. To resolve these cell-type–specific contributions, we analyzed bioenergetic gene expression in *Nkx2.1*-positive interneuron precursor populations^51^ using a publicly available single-cell RNA sequencing dataset with *Grin1* perturbation. Glycolytic gene expression was increased in hippocampal GABAergic neurons; an effect not seen in the cortical populations (Figure 5A). A similar pattern was observed for TCA cycle–related genes (Figure 5B), whereas genes associated with pyruvate metabolism and fatty acid oxidation showed no substantial changes in both populations (Figure 5C–D).

**Figure 5.**
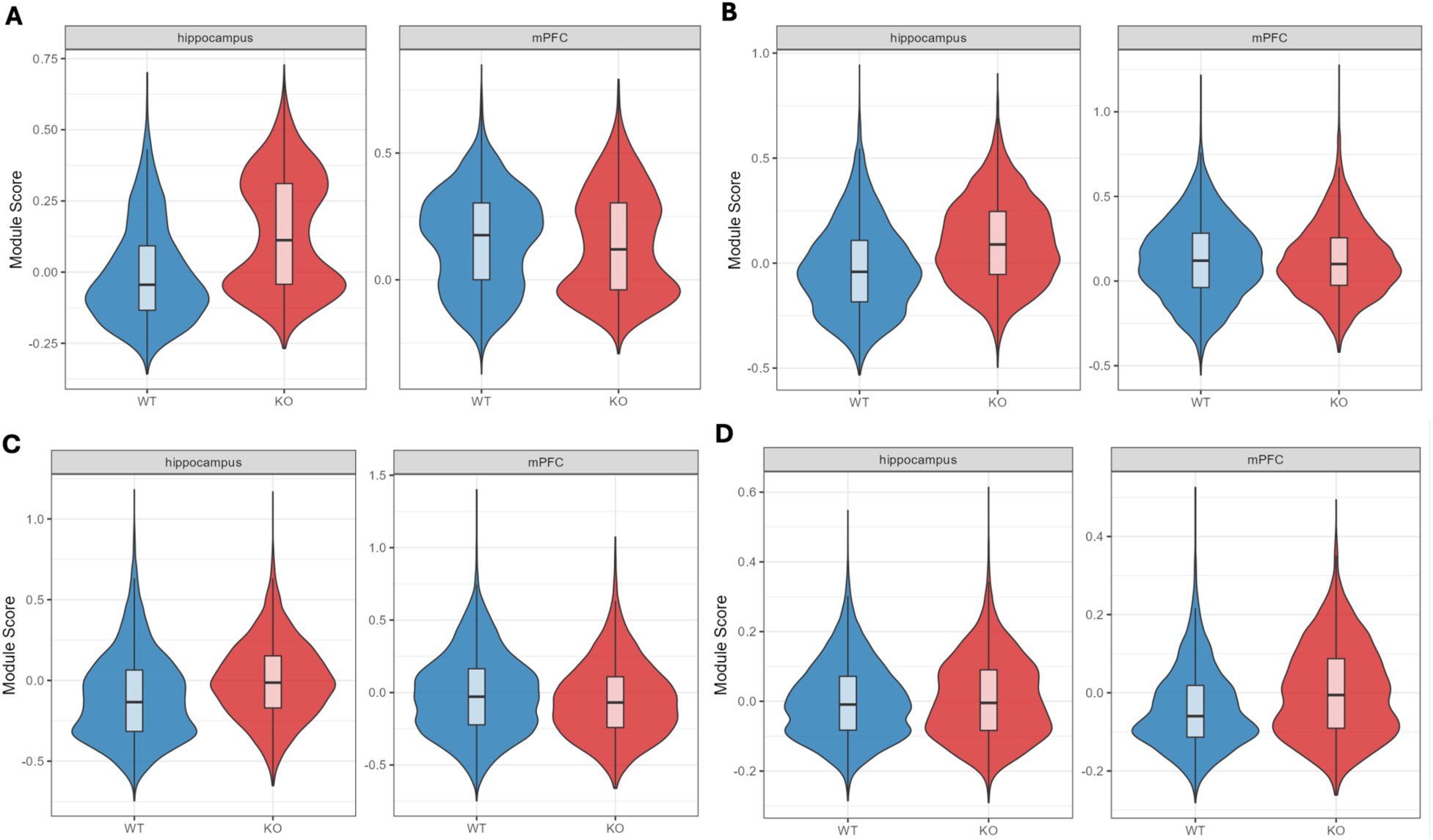
*Grin1* deficient interneurons show region specific bioenergetic profiles. Violin maps showing the expression pattern of bioenergetically relevant genes for (A) glycolysis, (B) pyruvate metabolism, (C) TCA cycle flux, and (D) fatty acid β-oxidation in *Nkx2.1:GRIN1*^fl/fl^ positive cells isolated from P18-21 hippocampus (left) and mPFC (right). Blue representing wildtype (WT) and red representing knockout (KO) mice, respectively.

## Discussion

### Early glycolytic transcriptional responses reveal neuroimmune–bioenergetic crosstalk following MIA

The earliest transcriptional alterations were detected 6 h post-MIA at E12.5, a stage dominated by radial glial cells, early neuronal progenitors, and microglia^52,53^. At this point, mature neuronal and glial populations are not yet established^54^, suggesting that the observed upregulation of glycolytic genes primarily reflects progenitor or non-neuronal metabolic activity. Interpretation therefore requires caution, as canonical adult cell-type bioenergetic signatures are not yet defined during early corticogenesis.

Consistent with prior reports, analysis of microglial markers revealed no coherent transcriptional evidence for increased abundance or classical activation states following MIA at E12.5. Based on our findings that Zika virus exposure, despite engaging TLR3-associated antiviral pathways, was insufficient to robustly induce native immune programs in early cortical progenitors, one could speculate that bulk transcriptomic analyses at this developmental stage are strongly influenced by the predominance of neural progenitor populations with intrinsically limited innate immune responsiveness. In this context, inflammatory signals arising from comparatively sparse immune-competent or more developmentally mature cell populations may become diluted within the broader transcriptional landscape, thereby reducing the detectability of classical inflammatory signatures.

Likewise, canonical inflammatory mediators were largely absent or below detection thresholds, paralleling observations by others^43,55^. Notably, *Trem2* showed a consistent directional increase across samples. Given its role in regulating microglial bioenergetics via the PI3K–AKT–mTOR pathway^56,57^, this trend may indicate a shift in microglial metabolic state independent of overt inflammatory activation. Together, these findings suggest that the observed transcriptional changes may be metabolically mediated, potentially through *Trem2*-associated pathways, rather than indicative of a classical immune responses, although this remains to be directly established.

Additional context comes from studies of early human cortical development, where neural progenitors exhibit limited innate immune responses^46^. Consistent with this, and with previous work^28^, ZIKV-exposed human cortical progenitors show minimal induction of classical inflammatory genes at early differentiation stages (radial glia and neural precursors), with responses increasing over time. Together, these findings support a model in which developmental stage and cell-type composition shape immune detection, such that early progenitors contribute little to classical inflammatory signatures.

Importantly, the distinction between global and cell-type–specific metabolic perturbations may have critical developmental consequences^58^. In humans, widespread bioenergetic disruption can result in severe outcomes, including fetal demise, whereas more restricted, cell-type-specific vulnerabilities may lead to selective anatomical or circuit-level alterations. For example, ZIKV preferentially targets cortical neuronal progenitors, resulting in microcephaly without uniformly affecting all cell populations^59^. In this context, the early glycolytic signature observed following MIA may reflect a) restricted or microglia-driven immune activity that is not readily detected in bulk transcriptomic analyses due to the predominance of less responsive cell populations or b) selective metabolic response that shapes downstream developmental trajectories.

### Developmental progression reveals dynamic glycolytic alterations

In contrast to E12.5, relatively few transcriptional changes were observed at E14.5, a period characterized by rapid neurogenesis and dynamic cortical remodeling^60^. As discussed by Molnár et al., 2012^53^, strong endogenous developmental programs related to progenitor proliferation, neuronal specification, and migration likely dominate the transcriptional landscape at this stage, potentially masking subtler environmentally induced perturbations. Nevertheless, early environmentally induced bioenergetic perturbations may have already primed these processes, shifting them onto an altered developmental trajectory that is not readily captured as overt transcriptional dysregulation at this stage. Consistent with this, increased *Pfkp* alongside reduced *Pdk3* suggests a subtle shift toward enhanced glycolytic flux with greater mitochondrial pyruvate utilization^29^. However, in the absence of coordinated regulation across related genes, this does not indicate a concerted pathway-level change but rather a modest signature compatible with a primed bioenergetic state.

By E17.5, when neuronal differentiation and laminar organization are more advanced^60^, glycolytic alterations in the neocortex of the MIA model re-emerged relative to vehicle controls. Isoform-specific analyses revealed increased expression of neuron-enriched genes alongside reduced expression of glial-associated transcripts. Together with changes in lactate metabolism, including increased *Ldha* and reduced *Ldhb*, these findings suggests that MIA promotes a shift towards an aerobic glycolytic state within neuronal populations. While aspects of this transcriptional profile resemble bioenergetic programs associated with early cortical expansion, differentiating neurons normally transition toward greater mitochondrial oxidative metabolism during this developmental window^12,61^, suggesting that the persistent glycolytic bias observed at E17.5 following MIA likely represents aberrant prolongation of immature metabolic programs. Alternatively, and not mutually exclusive, these changes may also reflect a compensatory adaptation to MIA-induced inflammatory stress.

Within this developmental context, additional transcriptional changes observed at E17.5 in pathways related to mitochondrial function and lipid metabolism suggest that glycolysis represents the dominant and most consistent metabolic signature in the MIA model, while mitochondrial and lipid programs likely act in a coordinated, supportive manner to sustain biosynthetic demands during late corticogenesis. This integrated metabolic configuration is consistent with research from Knobloch who postulated the role of mitochondria in proliferating and differentiating neural populations through to non-bioenergetic processes such as lipid synthesis for axonal growth and synaptogenesis^12^.

Importantly, the persistence of bioenergetic dysregulation in the MIA model may reflect the sustained influence of the maternal environment during gestation^43,55^. Unlike genetic perturbations, MIA represents a sustained, system-wide exposure in which maternal immune and metabolic alterations continuously influence the developing brain throughout gestation. In contrast, the absence of significant transcriptional changes in the *Srr⁻/⁻* model at this stage may reflect the more restricted nature of the perturbation, targeting a single metabolic–synaptic axis (D-serine/NMDA receptor co-agonism)^18,19^, which may be buffered by compensatory mechanisms^62^. Indeed, the extent to which glycine, an alternative NMDA receptor co-agonist, can compensate for reduced D-serine remains incompletely understood^63^ and may contribute to relative preservation of embryonic transcriptional programs in this model. Nevertheless, partial compensation at the transcriptional level may not fully restore activity-dependent developmental signaling or circuit maturation, potentially explaining how schizophrenia-like phenotypes can emerge despite comparatively modest bulk transcriptomic alterations^64,65^.

At this timepoint, focused on glycolytic transcription as the strongest upregulated process in the MIA model, we screened key genes in a genetic model (*Srr^-/-^*). Although qPCR revealed a consistent directional increase across most genes, no transcriptional upregulation was significant. This divergence likely reflects fundamental differences between environmental and genetic perturbations during late embryogenesis. MIA induces an acute, system-wide metabolic response, whereas loss of serine racemase primarily alters D-serine availability and NMDA receptor co-agonism, with downstream metabolic effects that may not yet be apparent at E17.5. Nevertheless, despite the lack of statistical significance in the *Srr⁻/⁻* model, the conserved directionality across both paradigms supports the presence of an early, sub-threshold bioenergetic adaptation rather than a robust, pathway-wide transcriptional shift.

Moreover, transcriptional glycolytic profiles at this stage could also be influenced by astrogliogenesis, adding nuance to the astrocyte–neuron lactate shuttle framework, as discussed by Kim et al., 2025^66^. Two non–mutually exclusive interpretations may explain this configuration. First, it may reflect delayed metabolic maturation, whereby neurons retain a more glycolytic state typically associated with immature populations^61^. Given the role of metabolic state in regulating neuronal differentiation, migration, and lineage specification^67^, early sub-threshold perturbations in the *Srr⁻/⁻*model may influence neurodevelopmental trajectories before becoming transcriptionally evident. Consistent with this, alterations in the ratios of proliferative neuronal populations, including phospho-histone H3–positive cells, have been reported in *Srr⁻/⁻* models^25^, supporting the idea that early bioenergetic disruptions may precede overt transcriptional phenotypes.

Second, the observed glycolytic upregulation may represent a distinct, developmentally regulated neuronal state that supports the high energetic and biosynthetic demands of late corticogenesis. In this context, increased expression of glycolytic enzymes, particularly *Hk2*, may indicate an active shift toward enhanced glycolytic flux. While *Hk2* is typically associated with glial metabolism, its induction in neurons has been reported under pathological conditions, including neurodegenerative states, where it promotes glycolysis and alters neuronal metabolic identity^68,69^. Accordingly, we proposed that *Hk2* upregulation in this model may reflect a neuron-intrinsic metabolic adaptation, potentially serving distinct functional roles across regions, rather than a purely glial-driven signal.

Together, these observations align with the concept of schizophrenia as a disorder of impaired metabolic flexibility^70^, in which neurodevelopmental systems may fail to appropriately transition between glycolytic and oxidative states during circuit maturation. This framework further supports the notion that the functional consequences of genetic risk may emerge more prominently at later developmental stages^71^, thereby motivating investigation during defined postnatal windows when compensatory mechanisms may no longer be sufficient.

### Juvenile bioenergetic specialization reveals region- and cell-type-specific vulnerability

Given the central role of cortical and hippocampal circuits in schizophrenia, juvenile period represents a critical window during which, particularly in the mPFC and hippocampus, neuronal metabolic states may influence synaptic maturation and circuit refinement; processes that are highly sensitive to NMDA receptor function and co-agonist availability^72^. In this context, increased expression of aerobic glycolytic genes in the mPFC of *Srr⁻/⁻* mice is consistent with ^31P-MRS studies reporting elevated high-energy phosphate metabolism in early-stage schizophrenia^5,73^. One possibility is that enhanced glycolysis supports not only energetic demands, but also biosynthetic pathways linked to serine metabolism, including D-serine production from glycolytic intermediates such as 3-phosphoglycerate^74^. This may represent a compensatory attempt to sustain NMDA receptor co-agonism under reduced serine racemase activity.

Upregulation of mitochondrial pyruvate transport and *Idh2* from the TCA flux screening in the mPFC suggests enhanced antioxidant capacity, consistent with the role of mitochondrial isocitrate dehydrogenase in NADPH production and redox homeostasis^75,76^. However, in the hippocampus, the elevation if *Idh2* accompanied by pyruvate efficient oxidation may instead reflect a form of metabolic reprogramming analogous to that observed in other high-demand metabolic states^77^. In addition, given the earlier maturation of the hippocampus relative to the mPFC^47^, this region-specific bioenergetic profile may reflect distinct temporal bioenergetic demands. Alternatively, region-specific bioenergetic profiles may reflect a disease-driven compensatory effect in which the hippocampus shift toward alternative substrate utilisation to preserve homeostatic brain nutrient processing; phenomenon that is consistent with reports showing reduced hippocampal ATP levels in individuals with schizophrenia^73^.

Interestingly, increased *Hk2* expression in both regions may not solely reflect glial activity. Although typically glial-associated, *Hk2* can be induced in neurons under pathological conditions, promoting glycolytic flux and altering neuronal metabolism^69^. In this context, elevated *Hk2* may represent a neuronal adaptation, or maladaptation, favouring glycolysis and underscoring the importance of cell-type-specific interpretation.

Determining whether this shift reflects a generalized feature of schizophrenia or a neuron-intrinsic metabolic phenotype requires cell-type–specific resolution. In this context, interneurons are of particular interest, given their well-established vulnerability in SZ^70,72^. To address this, we analyzed single-cell RNA sequencing data from Nkx2.1-derived interneurons with abolished *Grin1*. Despite convergence on NMDA receptor hypofunction, *Grin1* and *Srr⁻/⁻* models differ mechanistically^62^: *Grin1* loss disrupts receptor function directly, whereas *Srr* deficiency alters co-agonist availability, potentially preserving partial receptor activity. This distinction may underlie divergent metabolic adaptations. In *Grin1*-deficient interneurons, increased glycolytic and mitochondrial gene expression in the hippocampus may reflect a compensatory response to intrinsic synaptic inactivity, consistent with tight coupling between synaptic activity and energy metabolism^78^, whereas in *Srr⁻/⁻* mice, glycolytic shifts may be more closely linked to metabolic support of neurotransmitter and co-agonist synthesis. Notably, opposing patterns in cortical populations in *Grin1*-deficient interneurons further support region- and cell-type-specific vulnerability, consistent with recent transcriptomic evidence of divergent bioenergetic regulation across cortical microcircuits in schizophrenia^79^.

Taken together, these findings suggest that genetic risk factors for schizophrenia differentially shape bioenergetic states during the juvenile period, potentially reflecting both distinct molecular mechanisms and differences in developmental timing of NMDA receptor disruption ^80,81^. Accordingly, metabolic reprogramming may represent a convergent but mechanistically heterogeneous pathway across models.

In this context, MIA provided a framework to guide exploratory intrauterine timepoints, whereas *Nkx2.1:Grin1^fl/fl^* and *Srr^−/−^*enabled more targeted, pathway-specific interrogation. Nevertheless, broader transcriptomic profiling during intrauterine development in genetic models, alongside juvenile and region-specific analyses across environmental and multiple brain cell types of genetic models, will be necessary to fully capture bioenergetic phenotypes. Lastly, potential baseline differences in developmental state across *Nkx2.1:Grin1^fl/fl^*and *Srr^−/−^* models underscore the importance of incorporating maturation metrics to ensure accurate interpretation of comparative bioenergetic data^82^.

## Conclusion

In summary, our findings demonstrate that bioenergetic reprogramming is a developmentally dynamic, cell-type– and region-specific feature associated with schizophrenia risk. Following maternal immune activation (MIA), early embryonic glycolytic changes were primarily driven by radial glia and microglia, whereas neuronal populations in mid- to late gestation exhibited coordinated modulation of glycolysis and mitochondrial pyruvate metabolism. Importantly, this persistent glycolytic bias may alter the trajectory of cortical maturation and potentially prime circuit development, with consequences for later responsiveness to environmental stimuli. In the genetic *Srr^−/−^* model, intrauterine alterations were more restricted. During the juvenile period, region- and cell-type-specific adaptations became more pronounced, with the mPFC showing elevated glycolytic, mitochondrial pyruvate entry, and TCA-associated transcription, while hippocampal bioenergetic reprogramming weighted towards neuronal isoforms. In parallel, *Nkx2.1*-derived *Grin1*-deficient interneurons displayed region-dependent metabolic divergence, with hippocampal upregulation of glycolytic and TCA cycle pathways and opposing cortical responses, highlighting cell-type–restricted metabolic adaptations that may precede circuit-level dysfunction.

Collectively, support the concept that vulnerability to schizophrenia-relevant phenotypes may arise from temporally and regionally restricted windows of metabolic plasticity during development, when bioenergetic programs are most sensitive to immune, genetic, and activity-dependent perturbations. In MIA, this may reflect immune-driven metabolic adaptation, whereas in *Srr^−/−^*models it may relate to altered serine–glycine metabolism and compensatory support of NMDA receptor function, and in *Nkx2.1:Grin1^fl/fl^* perturbation may reflect activity-dependent energetic and migratory demands in interneuron populations. These divergent pathways converge on a common theme of altered metabolic flexibility across development, brain region, and cell type. Our work therefore highlights transcriptionally defined bioenergetic states as candidate contributors to neurodevelopmental vulnerability in schizophrenia and shared psychiatric disorders, warranting future functional validation.

## Strengths and limitations

This study leverages a multimodal approach to investigate schizophrenia-associated bioenergetic alterations, integrating both genetic (*Srr^−/−^*, *Grin1* ablation) and environmental (MIA) models across intrauterine and postnatal stages. By assessing bioenergetic profiles across intrauterine and postnatal stages, we captured temporal dynamics of metabolic changes from early corticogenesis to juvenile period, providing one of the first attempts to resolve developmental bioenergetic programs across multiple schizophrenia-relevant models using isoform- and cell-type–resolved approaches. By focusing on brain-specific isoforms and key metabolic pathways (glycolysis, mitochondrial function, and lipid oxidation), we provide a temporally resolved and mechanistically focused analysis of developmental bioenergetic changes across models. Nonetheless, several limitations should be acknowledged. Early intrauterine analyses in the MIA model relied on bulk RNA-seq with cell-type-specific resolution mostly limited to neurons versus glia, likely masking heterogeneity across glial subtypes and temporally distinct developmental populations. In addition, the *Srr^−/−^* intrauterine data were restricted to a single developmental timepoint and qPCR-based measurements rather than transcriptome-wide approaches, limiting molecular resolution and pathway-level inference. Juvenile *Grin1* analyses were limited to *Nkx2.1*-positive cells from a single animal, preventing statistical inference and reducing generalizability. In addition, the absence of juvenile MIA data constrains direct comparison between environmental and genetic perturbations at this stage. Finally, differences in baseline developmental states across models, particularly considering that genetic perturbations can alter regional maturation trajectories, highlight the importance of controlling for developmental stage when comparing bioenergetic phenotypes across paradigms. Future work incorporating transcriptomic profiling across additional developmental stages and cell types will be necessary to fully resolve convergent and divergent transcriptional bioenergetic programs in schizophrenia risk models. Also, future studies integrating enzymatic activity, metabolomic profiling, and functional assessments of mitochondrial and glycolytic flux across development will be essential to determine the extent to which these transcriptional programs translate into altered bioenergetic function. Such approaches will be critical to establish whether these developmental metabolic states actively drive circuit assembly or instead reflect compensatory adaptations to upstream perturbations.

## Acknowledgements

The authors would like to thank Dr. Catriona H. E. Rooney and Prof. William S. James for proofreading this manuscript. Special thanks go to Prof. Nikolaos P. Daskalakis and Prof. Kerry J. Ressler as well as additional members of the Metabolic and Mental Health Program, the Laboratory of Neurobiology of Fear and the Neurogenomics and Translational Bioinformatics Laboratory within the Department of Depression and Anxiety Disorders at McLean Hospital.

## Contributions

JG-J conceived and designed the study. OOF and ZM contributed to the study design. JG-J, OOF, MC-G and TVK collected and analyzed the data. OOF, HW, CMP and ZS contributed to the interpretation of the findings. JG-J drafted the manuscript, and all co-authors provided critical suggestions, read and accepted the final version prior to submission.

## Funding

This study was supported by philanthropic donations to CMP under the Metabolic and Mental Health program at McLean Hospital.

## Ethics declarations

All methods were performed in accordance with the relevant guidelines and regulations. The study analyzed publicly available datasets derived from previous studies in which ethical approval was acquired. For our model, Institutional Review Board approval was given in compliance with existing regulations for animal work.

---

**Supplementary table 1.**
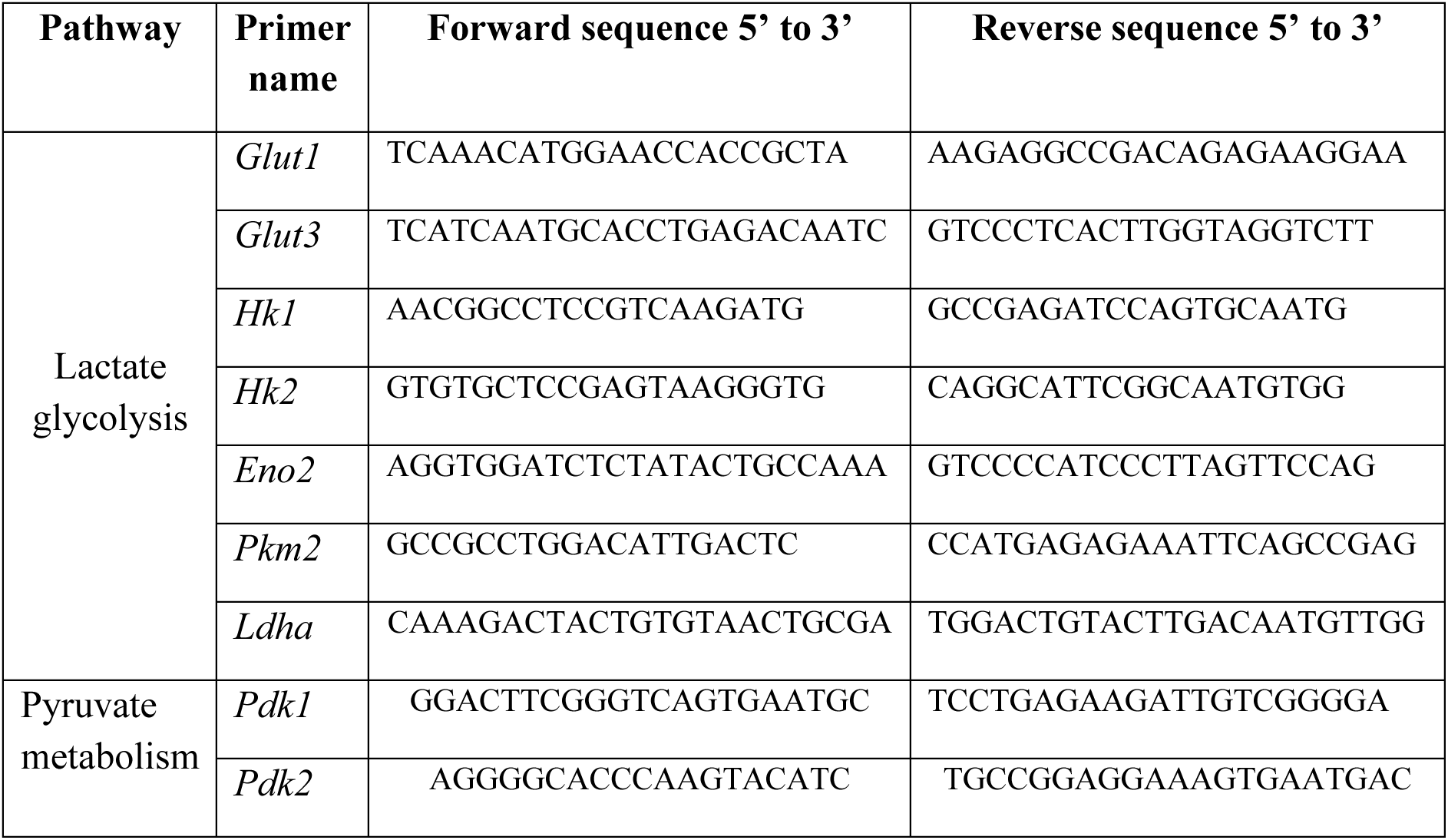
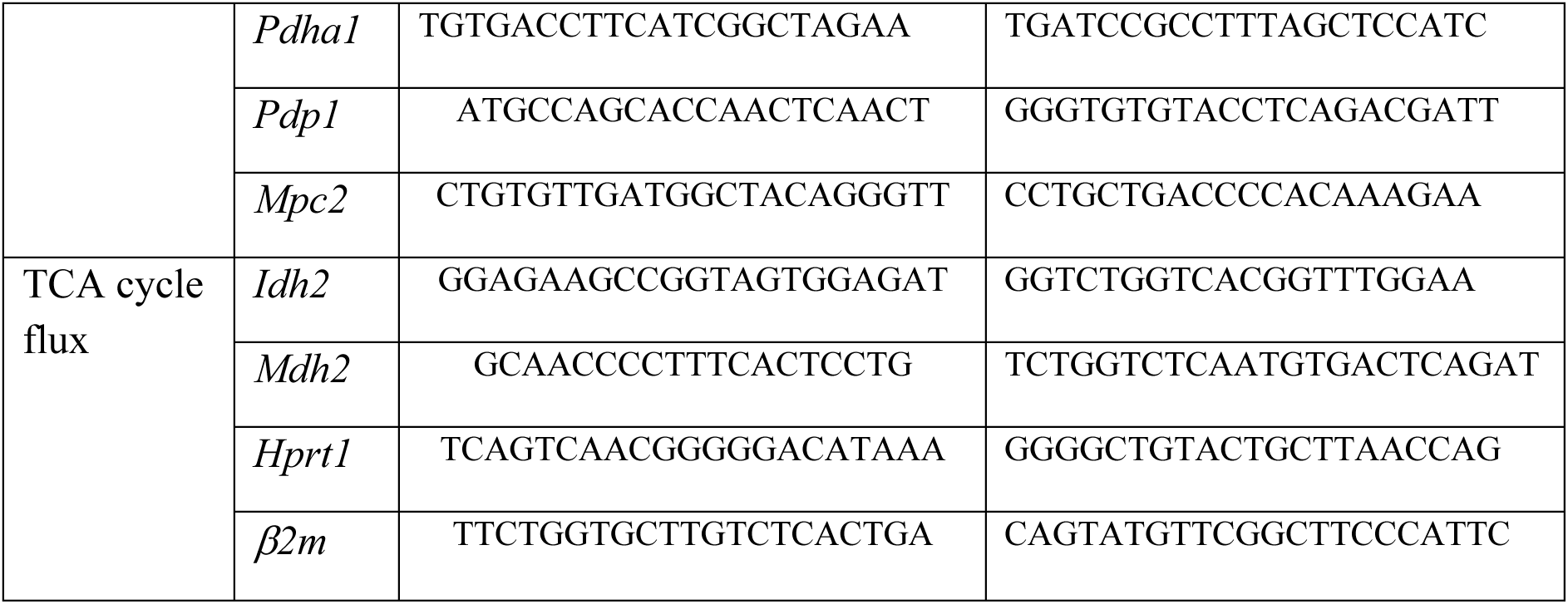
Primers sequences used for RT-qPCR analysis. Table containing all the primer pairs used for quantitative real-time PCR (RT–qPCR) to trace bioenergetic profiles in neocortex, juvenile cortex and juvenile hippocampus samples. Specificity was confirmed by melt curve assessment.

**Supplementary table 2.**
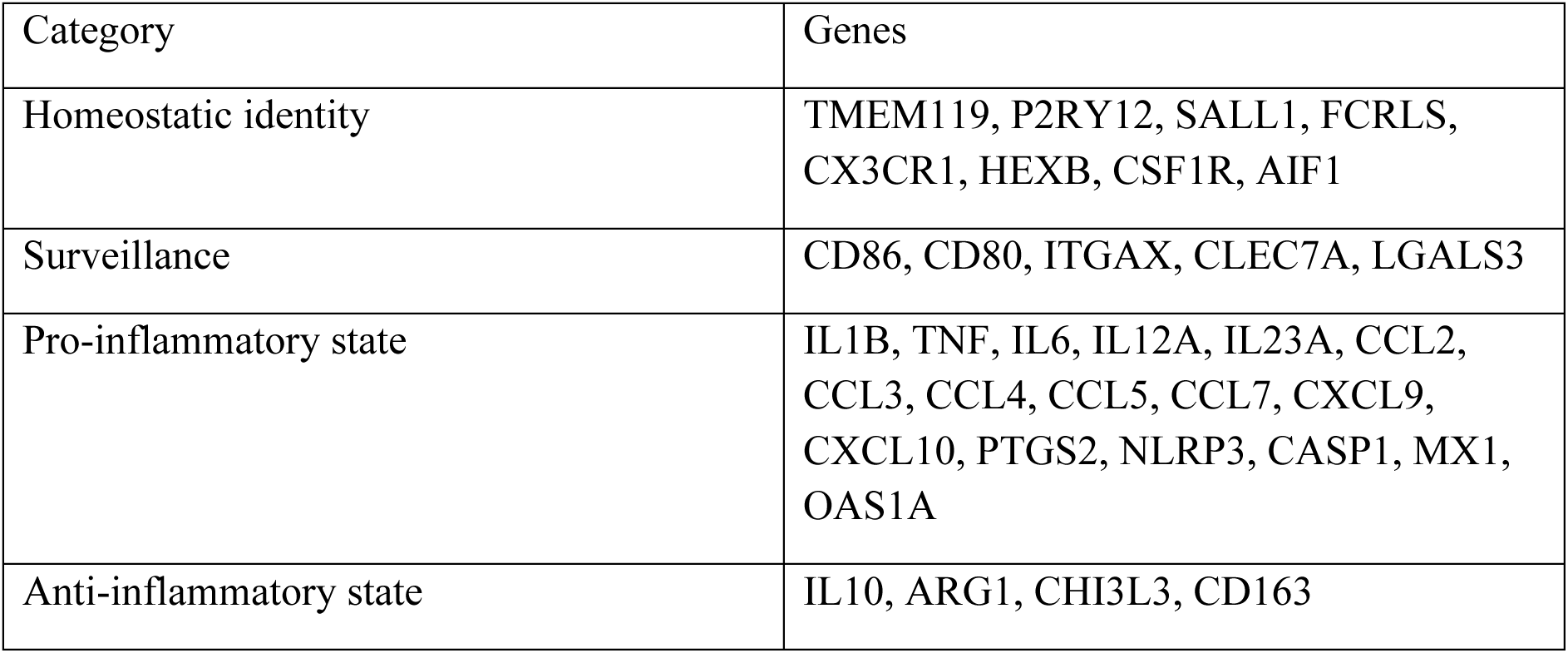
Microglial gene sets in MIA E12.5 neocortex. Gene sets corresponding to microglial categories that were curated based on the literature. The table lists specific genes that were not detected or not present in transcriptomic datasets for the MIA model at E12.5.

| Category | Genes |
| --- | --- |
| Homeostatic identity | TMEM119, P2RY12, SALL1, FCRLS, CX3CR1, HEXB, CSF1R, AIF1 |
| Surveillance | CD86, CD80, ITGAX, CLEC7A, LGALS3 |
| Pro-inflammatory state | IL1B, TNF, IL6, IL12A, IL23A, CCL2, CCL3, CCL4, CCL5, CCL7, CXCL9, CXCL10, PTGS2, NLRP3, CASP1, MX1, OAS1A |
| Anti-inflammatory state | IL10, ARG1, CHI3L3, CD163 |

**Supplementary Figure 1.**
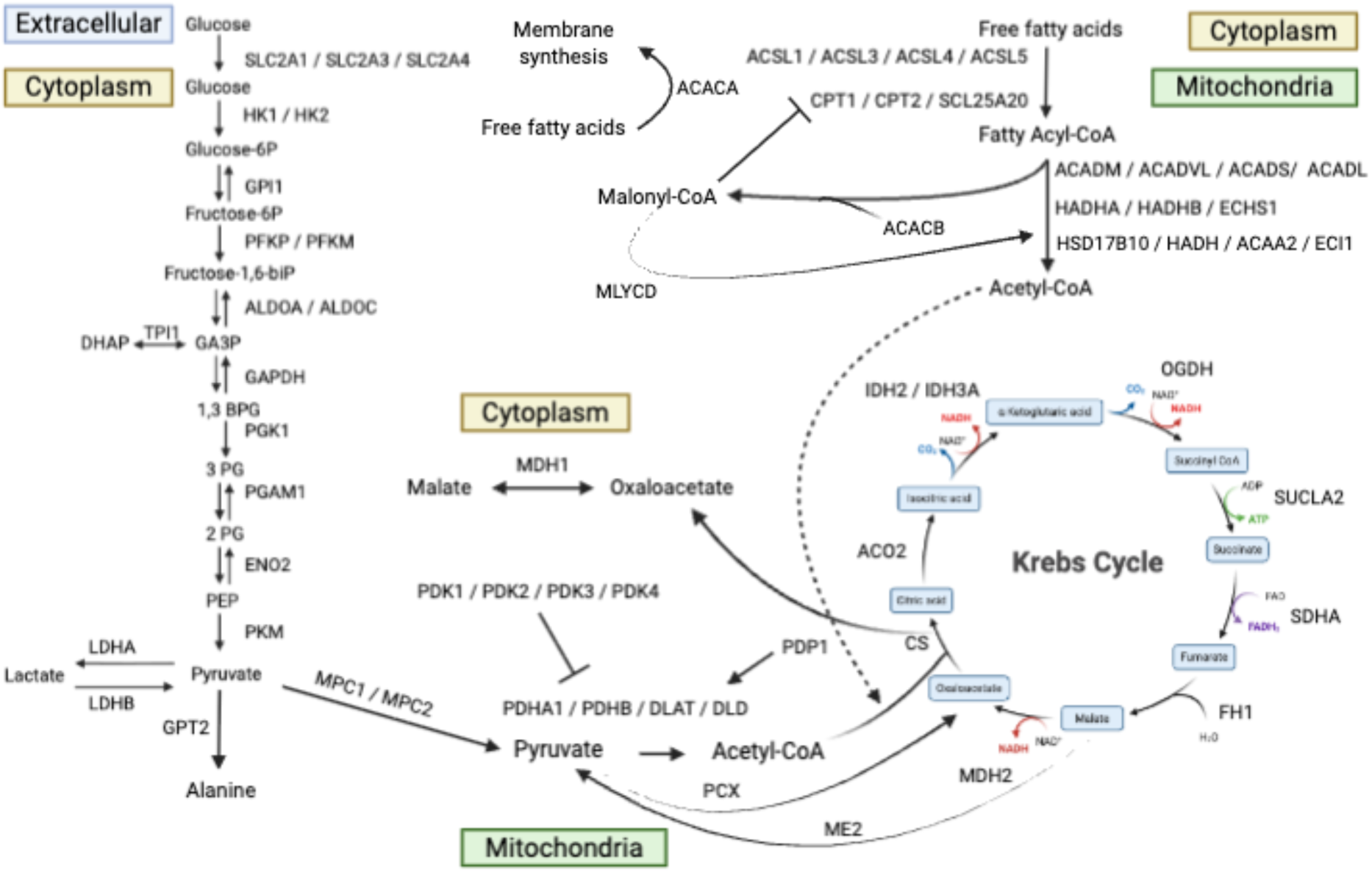
Integrated bioenergetic pathway map. Schematic representation of bioenergetic pathways integrating genes analysed from bulk RNA sequencing, single-cell RNA sequencing (scRNA-seq), and RT–qPCR datasets. Genes are organized within pathway-specific modules including glycolysis, lactate and pyruvate metabolism, tricarboxylic acid (TCA) cycle, and fatty acid β-oxidation. Arrows indicate the directionality of biochemical reactions or regulatory influence. Single-headed arrows denote unidirectional processes, whereas bidirectional arrows represent reversible reactions. Converging and branching arrows highlight points of metabolic crosstalk and pathway integration. Dotted lines indicate equivalent metabolites or intermediates generated from distinct pathways that could not be spatially represented within the same branch of the schematic. Cellular compartmentalization is indicated in colored boxes, distinguishing processes occurring in the cytosol, mitochondria, and extracellular space.

**Supplementary Figure 2.**
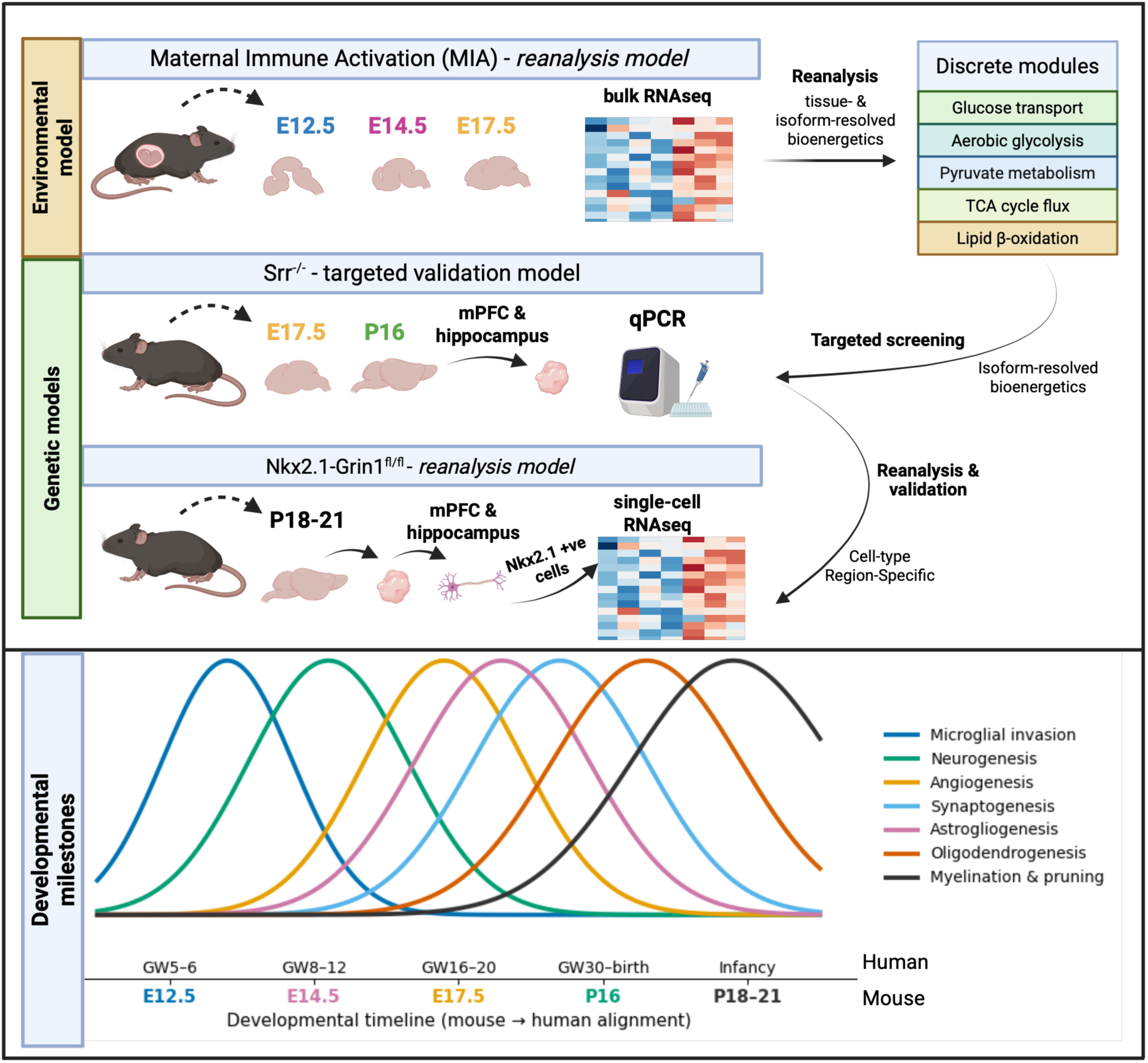
Experimental design and integrative analytical framework across developmental stages. Schematic overview of the experimental design across the three schizophrenia risk models used in this project. Each diagram illustrates developmental stages, tissue collection timepoints, and analytical approaches. For each model, each diagram illustrates tissue collection timepoints, analytical approaches, and type of tissue collected. Analytical modalities include RT–qPCR and re-analysis of publicly available bulk RNA-seq and single-cell RNA-seq (scRNA-seq) datasets. Genes were grouped into discrete bioenergetic pathway modules. Bottom panel representing a developmental timeline with color-coded bell-shaped curves representing key neurodevelopmental processes, highlighting periods of peak activity and relative temporal overlap across mice and humans.

**Supplementary Figure 3.**
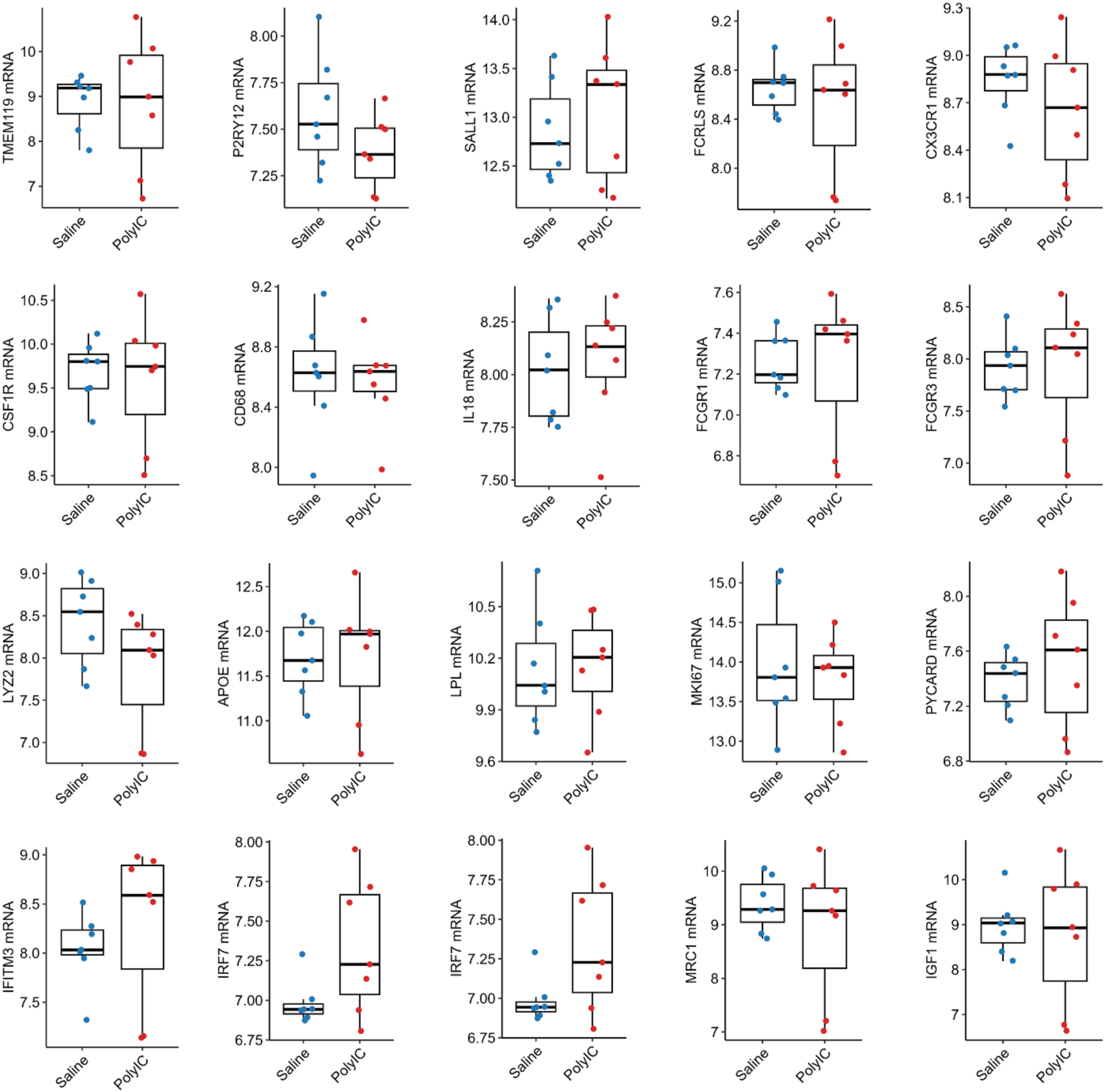
Microglial gene expression in the MIA model at E12.5. Box plots showing mRNA expression levels of microglia-associated genes at E12.5 in the MIA model. Red and blue dots represent MIA and saline control groups, respectively. Box plots show median (interquartile range). Statistical significance (adjusted p-values) was determined using DESeq2. No statistically significant differences were observed.

**Supplementary Figure 4.**
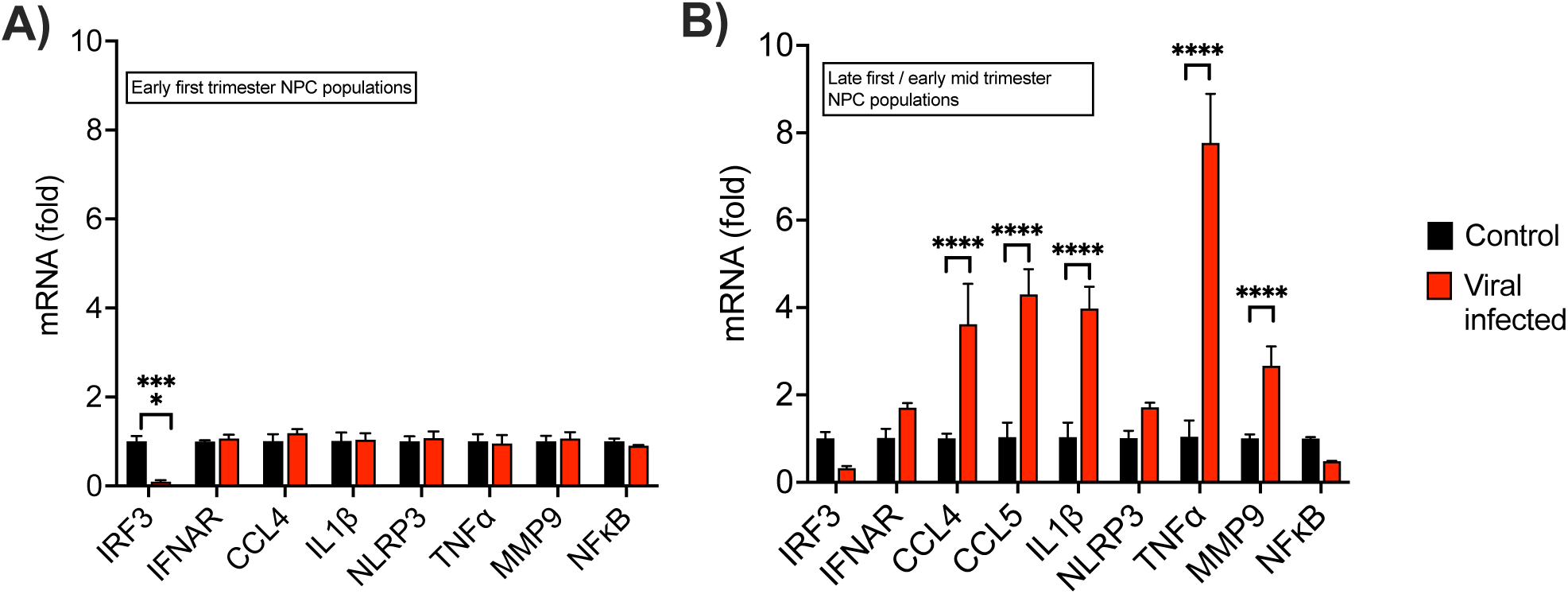
Innate immune gene expression in ZIKV-challenged iPSC-derived cortical progenitors. Bar graphs showing mRNA expression levels of innate immune-related genes in iPSC-derived cortical progenitors at two stages of differentiation: (A) early and (B) late stages, corresponding to early first trimester and late first/early mid-trimester neurodevelopmental timepoints, respectively. Cells were treated with Zika virus (ZIKV) for 24 h at an M.O.I. of 1. Black and red bars represent control and ZIKV-challenged conditions, respectively. Statistical analysis was performed using a two-way ANOVA followed by post hoc comparisons with Sidak’s correction. Data represent three independent cell lines and are shown as mean ± SD. Significance is indicated as * p ≤ 0.05, ** p ≤ 0.005, and *** p ≤ 0.0005.

## Notes

### Competing Interest Statement

The authors have declared no competing interest.

